# Phenotypic profiling, Comparative analysis of Nutritional compositions and Antioxidant potential of *Pleurotus ostreatus* (oyster mushroom) cultivated on Different Agricultural wastes

**DOI:** 10.1101/2025.08.09.669472

**Authors:** Olagunju Johnson Adetuwo

## Abstract

Oyster mushrooms (Pleurotus ostreatus) are a type of edible mushroom that has gained popularity worldwide due to its nutritional benefits, culinary value, and potential medicinal properties. This study investigated the phenotypic profiling, nutritional compositions, and antioxidant potential of Pleurotus ostreatus (oyster mushroom) cultivated on three different agricultural wastes: Gmelina tree sawdust, oil palm fruit pressed fibers, and cassava peels using standard mycological procedures. The results showed significant variations in morphological features, nutritional content, and bioactive compounds among the mushrooms grown on different substrates. Pleurotus ostreatus grown on oil palm fruit pressed fibers looked blossomed, tan in color, and matured faster, while those grown on cassava peels did not blossom, had a shiny white color, and matured late. *Pleurotus ostreatus* grown on Gmelina tree sawdust had the highest protein content (9.8%), while those on oil palm fruit fibers had the highest fiber content (6.6%) and steroid content (75.6%). Mushroom cultivated on cassava peels exhibited the highest carbohydrate content (13.3%), saponins-like compounds (39.6%), and antioxidant activity (RSA: 74.9%, IC50: 2.4 mg/ml). These findings suggest that the substrate used for cultivation significantly influences the nutritional and bioactive properties of *P. ostreatus*, allowing for the production of mushrooms with specific desired characteristics.

## 1.0. Background of the study

Oyster mushrooms (Pleurotus ostreatus) are a type of edible fungi that have gained popularity worldwide due to their nutritional benefits, culinary value, and potential medicinal properties (Degreef and de-Kesel, 2022; Fakoya, 2025) These mushrooms are rich in protein, fiber, and various vitamins and minerals, making them an excellent addition to a healthy diet. Moreover, oyster mushrooms have been shown to possess antioxidant, anti-inflammatory, and antimicrobial properties, which can contribute to overall well-being (Fakoya, 2025).

However, the increasing demand for food production has led to a significant amount of agricultural waste generation. In Nigeria, for example, agricultural waste management remains a significant challenge, with many farmers resorting to open burning or disposal in water bodies. This not only contributes to environmental pollution but also wastes valuable resources.

This study aims to explore the potential of utilizing agricultural waste as a substrate for oyster mushroom cultivation. By investigating the phenotypic profiling, nutritional composition, and antioxidant potential of oyster mushrooms grown on different agricultural wastes, this research seeks to contribute to sustainable agriculture, improve food security, and enhance nutrition.

The significance of this study lies in its potential to promote sustainable agricultural practices, reduce waste management issues, and provide a valuable source of nutritious food. The findings of this research can also inform policy decisions and agricultural practices, ultimately contributing to a more sustainable and food-secure future.

## 2.0 Literature Review

### 2.1. Oyster Mushroom Cultivation

Oyster mushrooms are one of the most widely cultivated edible mushrooms globally. They can be grown on a variety of substrates, including agricultural waste, forestry waste, and industrial waste. The cultivation of oyster mushrooms on agricultural waste has been shown to be a viable option for waste management and sustainable agriculture (Akinrinola-Akinyemi et al., 2017; Ojha et al., 2025).

Studies have demonstrated that oyster mushrooms can be successfully grown on substrates such as straw, sawdust, and other lignocellulosic materials. The use of agricultural waste as a substrate for oyster mushroom cultivation can help reduce waste management issues, promote sustainable agriculture, and provide a valuable source of income for farmers (Mubashar et al., 2024).

### 2.2. Nutritional Composition of Oyster Mushrooms

Oyster mushrooms are a nutrient-rich food, high in protein, fiber, and various vitamins and minerals (Fakoya et al., 2014). They are an excellent source of essential amino acids, including glutamic acid, aspartic acid, and arginine. Oyster mushrooms are also rich in minerals such as potassium, phosphorus, and copper.

The nutritional composition of oyster mushrooms can vary depending on the substrate used for cultivation. Studies have shown that oyster mushrooms grown on agricultural waste substrates can have higher nutritional content compared to those grown on traditional substrates (Ogundele et al., 2017).

### 2.3. Antioxidant Potential of Oyster Mushrooms

Oyster mushrooms have been shown to possess antioxidant properties, which can contribute to their potential health benefits (Alves et al., 2013; Ade-Ogunnowo et al., 2024). The antioxidant activity of oyster mushrooms is attributed to the presence of bioactive compounds such as phenolic acids, flavonoids, and polysaccharides (Kosti’c et al., 2017; Bakir et al., 2018).

Studies have demonstrated that oyster mushroom extracts can scavenge free radicals, reduce oxidative stress, and protect against cell damage. The antioxidant potential of oyster mushrooms can vary depending on the substrate used for cultivation, with some studies suggesting that mushrooms grown on agricultural waste substrates may have higher antioxidant activity (Chang et al., 2018; Koch, 2020).

### 2.4. Agricultural Waste Management

Agricultural waste management remains a significant challenge globally, with many farmers resorting to open burning or disposal in water bodies. This not only contributes to environmental pollution but also wastes valuable resources.

The use of agricultural waste as a substrate for oyster mushroom cultivation can help reduce waste management issues and promote sustainable agriculture. By utilizing agricultural waste, farmers can generate income, reduce environmental pollution, and promote sustainable agricultural practices.

### 2.5. Sustainable Agriculture and Food Security

Sustainable agriculture and food security are critical issues globally, with the world’s population projected to reach 9.7 billion by 2050 (FAO, 2024). The use of agricultural waste as a substrate for oyster mushroom cultivation can contribute to sustainable agriculture and food security by promoting efficient use of resources, reducing waste management issues, and providing a valuable source of nutritious food.

By adopting sustainable agricultural practices, farmers can improve crop yields, reduce environmental pollution, and promote food security. The cultivation of oyster mushrooms on agricultural waste can be a valuable strategy for promoting sustainable agriculture and improving food security, particularly in regions with limited access to nutritious food (Valverde et al., 2015).

## 3.0 Materials and Methods

### 3.1. Collection of Pleurotus ostreatus

In the course of this study, the strain of Pleurotus ostreatus was carefully sourced from a collection of preserved spawn mycelium in microbiology laboratory at the Department of Biological Sciences at Olusegun Agagu University of Science and Technology. This institution, located in the vibrant town of Okitipupa in Ondo State, Nigeria. The spawn mycelium was re-cultured onto Potato Dextrose Agar for 72 hours before it was inoculated onto bulky substrates (Gmelina sawdust, Cassava peels and Oil palm fruit fibers respectively)

### 3.2. Cultivation of Pleurotus ostreatus

In this study, the cultivation of the Pleurotus ostreatus was done using different agricultural wastes as the substrates, specifically Gmelina arborea tree sawdust, oil palm fruit fibers, and cassava peels. These materials were selected due to their widespread availability and potential as sustainable substrates for mushroom cultivation. The cultivation process adhered closely to established standard methodologies designed for optimizing the growth conditions of Pleurotus ostreatus (Prodhan et al., 2015; Mubashar et al., 2024). The selected sawdust, fibers, and peels were prepared through a series of preprocessing steps, which included thorough cleaning, drying, roughly grinding (for cassava peels) and subsequent sterilization (autoclaved at 1210C) to eliminate any microbial contaminants that could interfere with fungal growth (Ade-Ogunnowo et al., 2024). The substrates were then inoculated with a selected strain of Pleurotus ostreatus in a controlled environment: The optimum conditions adopted were: Temperature (25°C (77°F), Humidity (85% relative humidity) and low light intensity (under shade). As direct sunlight can be detrimental. Maintaining optimal conditions ensured healthy growth and high yields (Pereira et al., 2013).

### 3.3. Processing of the mushroom samples

Once the mushrooms had matured, a harvesting process was carried out, wherein the fruiting bodies were carefully collected to prevent any damage to the mycelial network. Following collection, the harvested Pleurotus ostreatus mushrooms underwent a drying phase utilizing a conventional drying method to preserve their structural integrity and nutrient profile. This drying process was essential in ensuring that the moisture content of the mushrooms was sufficiently reduced, thereby extending their shelf life and enhancing their viability for subsequent analyses (AOAC, 2020; Ade-Ogunnowo et al., 2024).

### 3.4. Extract Preparation

Mushroom extracts were made by drying whole mushrooms and then grinding them into a fine powder. The powder from each sample of mushroom was then dissolved in ethanol and stirred (20 v/v; 30 ml) at 25 °C and 150 rpm for 1 h, followed by filtration using Whatman filter paper number 2. The two combined fractions were then evaporated using a rotary evaporator at 40 °C under reduced pressure, allowing the ethanolic portion of the extract to evaporate. The resulting aqueous extract was refrigerated at 4 °C and stored for further use.

### 3.5. Comparative analysis of nutritional compositions of the mushroom samples

Subsequently, a comprehensive nutritional analysis was conducted to quantify the moisture content, protein, fiber, carbohydrates, fatty acids and mineral content of the dried mushrooms. This analysis employed standard laboratory techniques including proximate analysis and spectrophotometry to provide accurate and reliable measurements of these major nutritional components (AOAC, 2020).

### 3.6. Comparative analysis of the bioactive compounds present in the mushroom samples

The bioactive compounds present in Pleurotus ostreatus were determined. These analyses were conducted using high-performance liquid chromatography (HPLC) and UV-Vis Spectroscopy to determine the vitamins content, the presence and concentration of phenolic acids, steriods content, saponins-like compounds and tannins content (Ribeiro et al., 2008; AOAC, 2020; Ade-Ogunnowo et al., 2024), which are known for their potential health benefits.

### 3.7. Evaluation of antioxidant activity in Pleurotus ostreatus cultivated on different agricultural wastes

The antioxidant activity of mushrooms samples was evaluated by a non-cell-based procedure, used a spectrophotometric analysis based on the scavenging of free radicals, using the 1-1-diphenyl-2-picrylhydrazyl (DPPH) radical, where the decrease in absorbance at 517nm indicates the antioxidant potential of each mushroom extract (Baliyan et al., 2022).

The procedures for DPPH assay used in this study are outlined below:

Two centimeter of mushroom extract was mixed with a 2 cm of DPPH solution in 2 cm methanol. The mixtures were incubated for 30 minutes in the dark. Then, the absorbance of the mixture was measured at 517nm with spectrophotometer. The higher antioxidant potential is indicated by decreasing in absorbance below 517nm. Then, the RSA (%) was determined by using the formula of Baliyan et al. (2022) as given below.

RSA (%) = [(Ac-As)]/Ac x100. Where, As =the absorbance of the sample and Ac=the absorbance of the negative control (DPPH solution). The values obtained were compared with a standard curve generated using ascorbic acid. Finally, the results were expressed as the percentage of DPPH radical scavenging activity.

### 3.8. Data Analysis

Three samples of P. ostreatus were analyzed for this study, and the antioxidant assays was carried out in triplicate. The results were expressed as mean values. The differences between samples were analyzed using one-way analysis of variance (ANOVA). This analysis was carried out using the SPSS v.22.0 software. The Principal Component Analysis (PCA) was performed to evaluate if chemical and functional characteristics could separate the samples analyzed. The PCA was applied to standardized variables (centered by the mean and scaled by the variance).

## 4.1. Results

The results showed that the substrate used for cultivation significantly affected the morphological features, the maturity rate, the nutritional content and bioactive compounds of the mushrooms. *Pleurotus ostreatus* cultivated on Gmelina and oil palm fruit pressed fibres grown and matured faster than the one cultivated on cassava peels as shown in table 1. It was also revealed in table 1 and plates 1, 2 and 3 that the colour of mushroom varied from white to light-brown based on the substrate in which they were cultivated, the *Pleurotus ostreatus* grown on Gmelina sawdust and cassava peels are whitish in colour while those ones that grown on oil palm fruit fibres had brownish to dull-white in colour. Again, it was observed in table 1 that the sizes and weights of fruiting bodies varied based on the substrate in which they were grown, the *P. ostreatus* grown on oil palm fibres had highest of 3.8 g. In relation to their nutritional compositions, the *Pleurotus ostreatus* cultivated on Gmelina tree sawdust had higher protein content as shown in table 2 (9.7%), while those cultivated on oil palm fruit fibres had higher fibre content, (6.6%). Also, table 2 showed that the *P. ostreatus* grown on cassava peels had higher carbohydrates content (13.3%). *Pleurotus ostreatus* had varying levels of sodium, potassium, calcium, Iron and zinc based on the substrate used for its cultivation, with the highest percentage observed in mushroom grown on Gmelina sawdust, closely followed by *P. ostreatus* grown on oil palm fruit fibers, as shown in table 2. Sodium content was high in *P. ostreatus* grown on Gmelina sawdust and oil palm fruit fibers, respectively, but low in *Pleurotus ostreatus* cultivated on cassava peels (1.8%). Potassium content was very high for the *P. ostreatus* species (1.4%, 1.6% and 2.0 %), and calcium contents were low (0.8%, 0.9%, and 0.8%). Also the vitamins content varied based on the substrate in which they grown, highest vitamins content was observed in *Pleurotus ostreatus* grown on oil palm fruit fibers (3.3 %). Furthermore, the agricultural wastes substrates used for their cultivation have effect on their shelf life as shown in table 1. The figure 1 revealed the presence of phenolic compounds, saponins, steroids, and tannins in the methanolic extracts of the selected mushroom species. From the graphical analysis, it shows that tannins and phenolic compounds contents were higher in *P. ostreatus* grown on Gmelina sawdust and cassava peels respectively but had lower steroids content. Also *P. ostreatus* grown on oil palm fruit fibers had the highest steroids content of 75.6% compared with ones grown on other substrates. The saponins-like compounds were higher in P. ostreatus grown on cassava peels substrate (39.6%), while lower content was analysed in *Pleurotus ostreatus* grown on Gmelina sawdust substrate (23.5%). The antioxidant potential of mushroom also varies based on the substrate used to cultivate it. The higher antioxidant potential was observed in *P. ostreatus* grown on cassava peels with antioxidant activity of RSA (74.9 %) and IC_50_ (2.4 mg/ml) while lower antioxidant activity was recorded in *P. ostreatus* extract cultivated on Gmelina sawdust (RSA 46% and IC_50_ 4.7 mg/ml).

**Plate 1.**
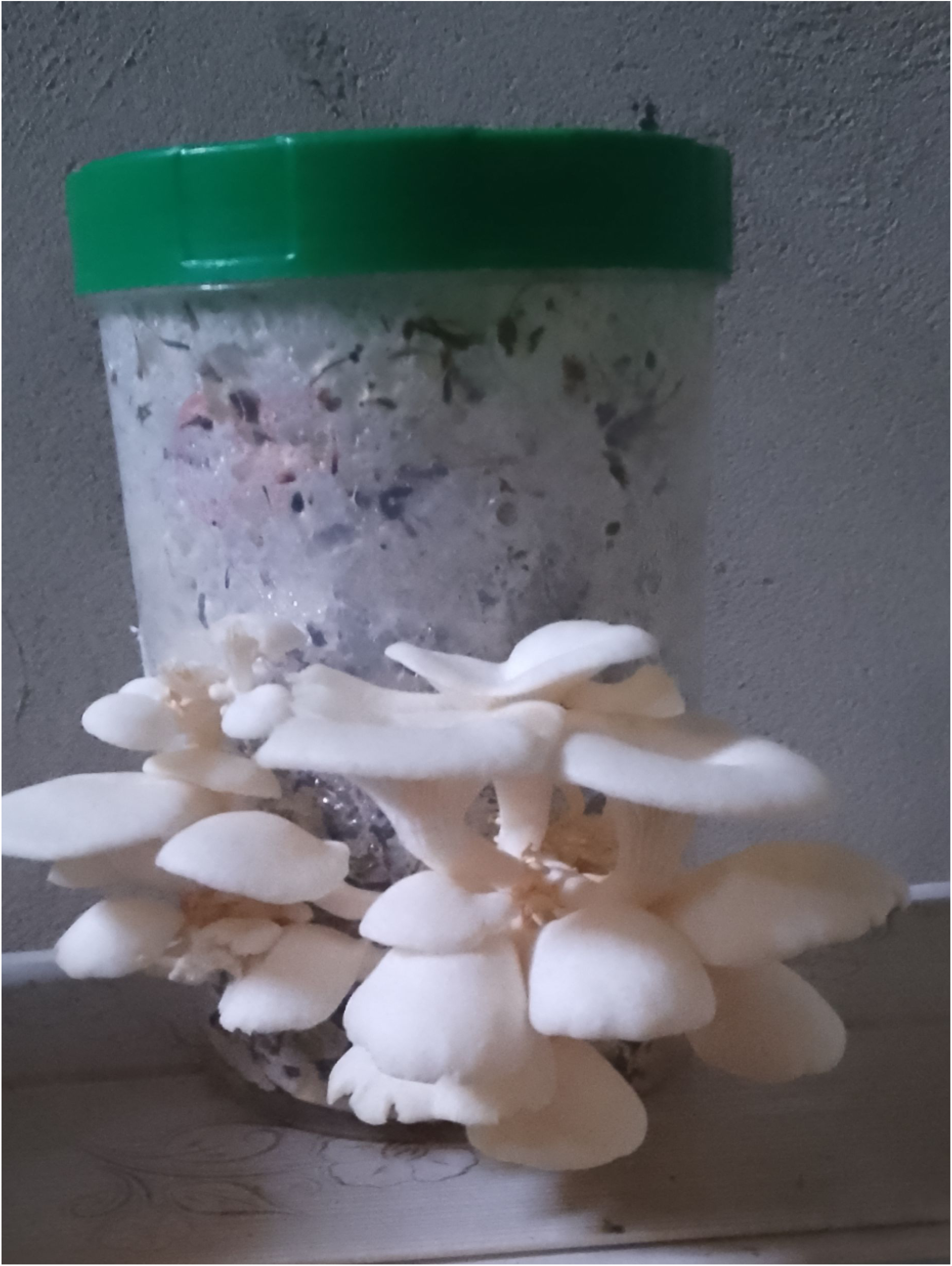
Pleurotus species cultivated on oil palm fruit pressed fibre

**Plate 2.**
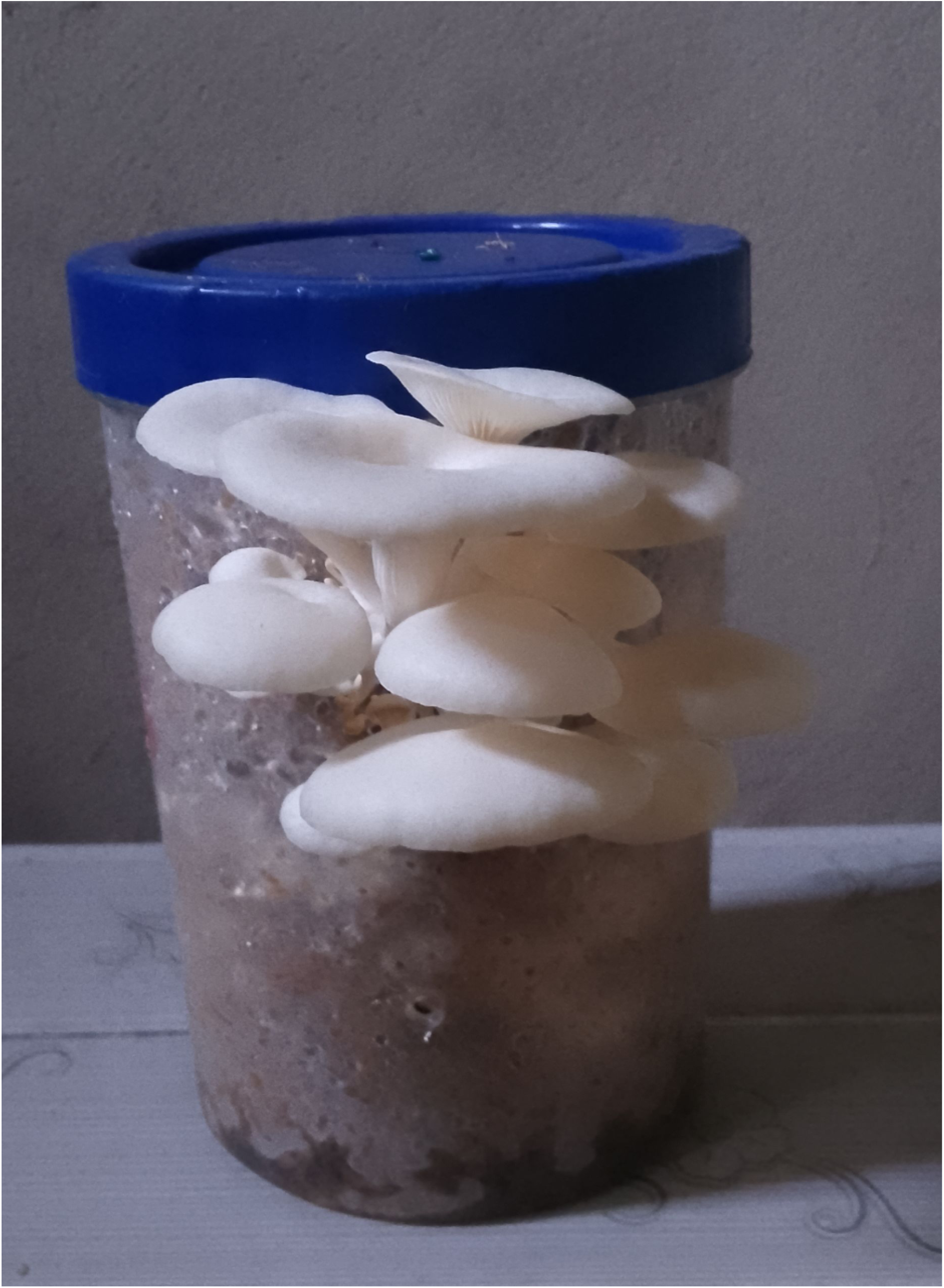
Pleurotus species cultivated on Gmelina sawdus

**Plate 3.**
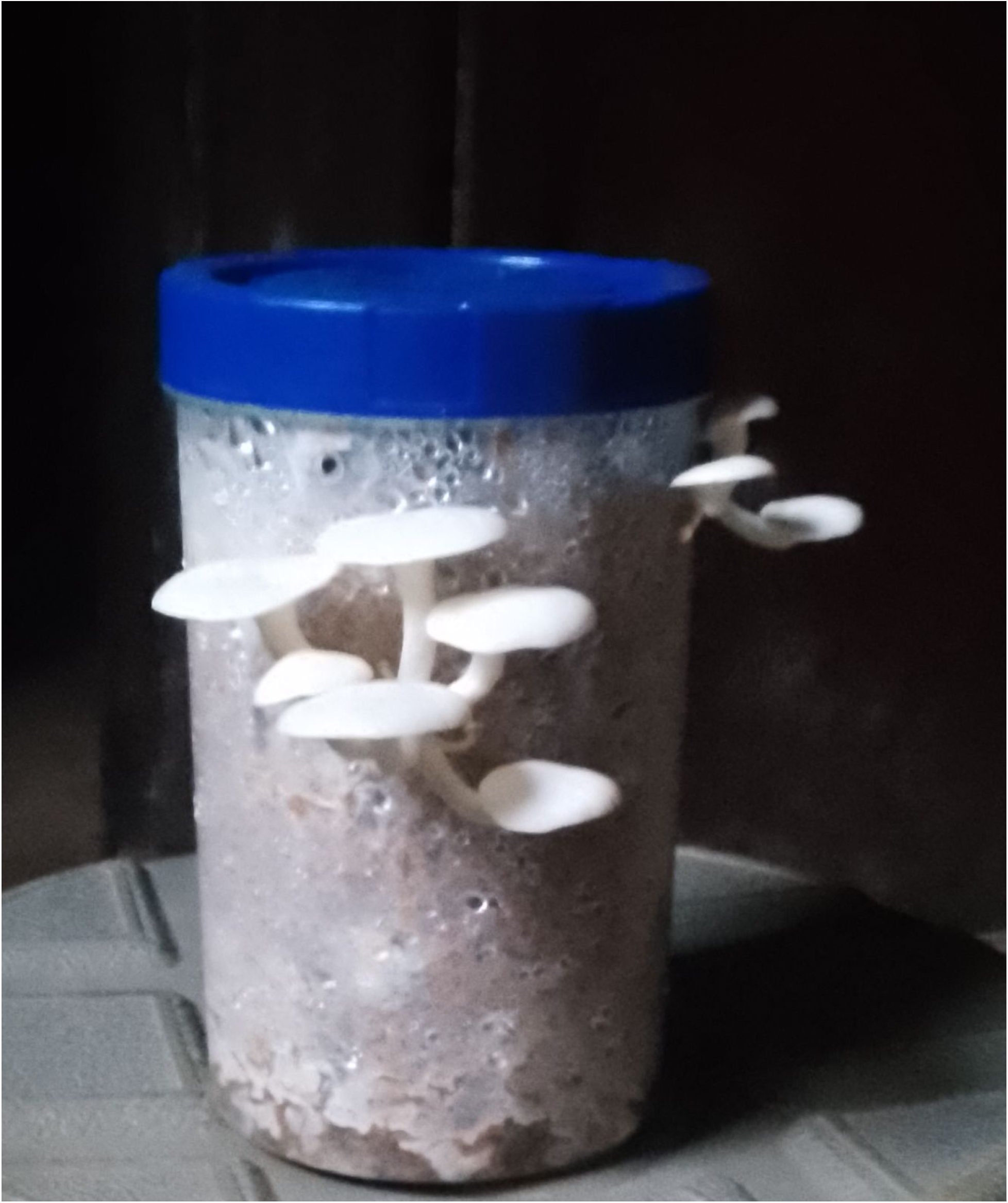
Pleurotus species cultivated on cassava peels

**Table 1.** Features of the fruiting of *Pleurotus species* cultivated on Gmelina sawdust, Cassava peels and Oil palm fruit pressed fibres respectively.

| Features | Gmelina sawdust | Cassava peels | Oil palm fruit fibres |
| --- | --- | --- | --- |
| Formation of pinhead in days after inoculation | 22 | 24 | 21 |
| Pinhead length in cm | 2.5 | 1.5 | 3.0 |
| Emergence of fruiting in days | 23 | 24 | 23 |
| Fruiting body height @ 22, 23 and 24 days in cm | 5.2 | 3.8 | 6.0 |
| Fruiting colour | White (dull) | White shiny | Tan |
| Pileus colour | White | White | Brownish-white |
| Pileus size in cm @ 25 day | 4.0 | 2.4 | 5.1 |
| Shape of pileus | Flat shape | Flat shape | Flat shape with curve edge |
| Presence of annulus | Lack | Lack | Lack |
| Colour of lamellae | White | Whit | Yellowish-white |
| Stipe length in cm | 4.3 | 3.1 | 4.6 |
| Mycelium growth rate in % | 30 | 22 | 35 |
| Fruiting body growth rate in % | 26 | 25 | 29 |
| Average shelf life | 36 hours | 48 hours | 30 hours |

**Table 2:**
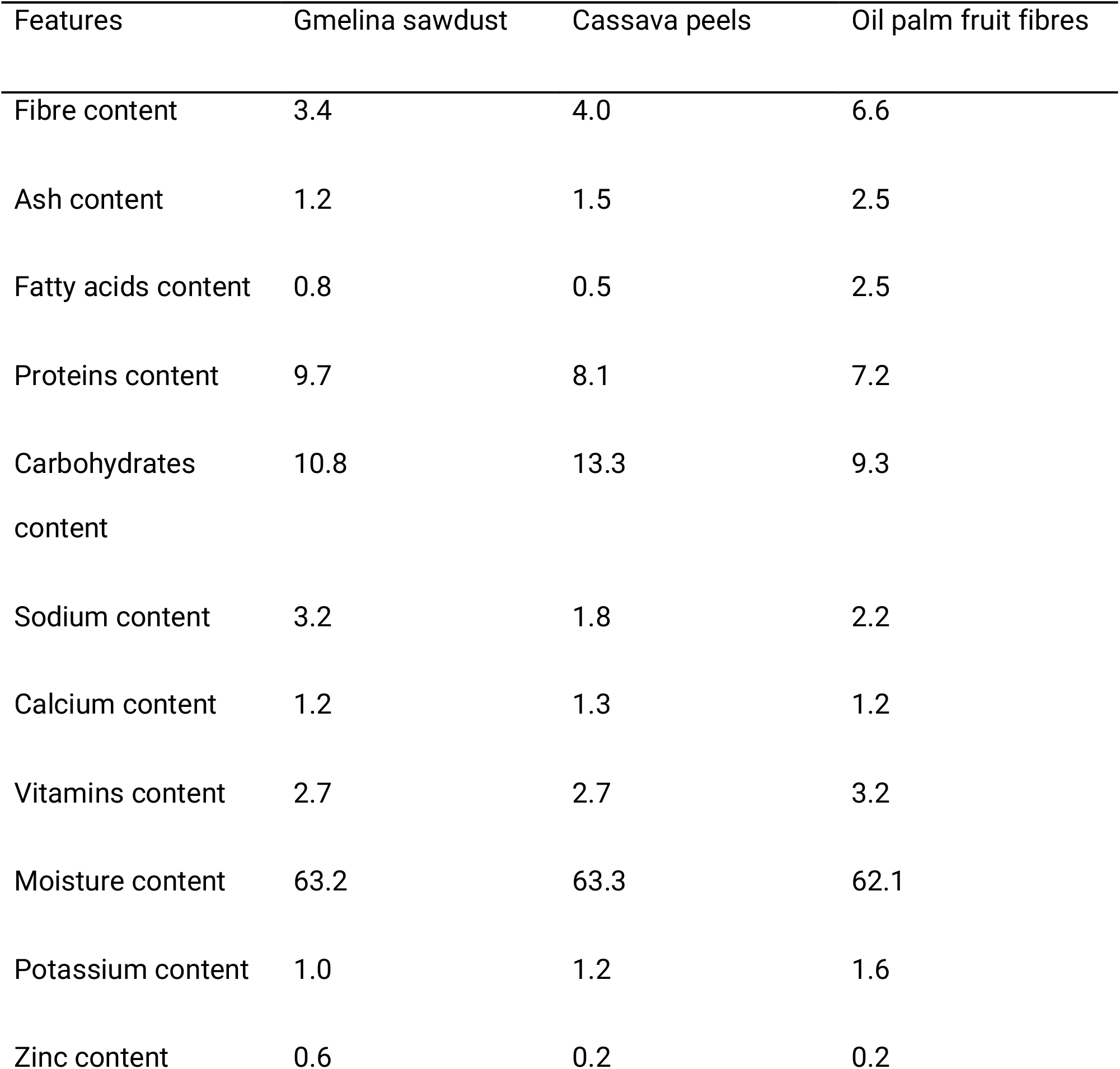
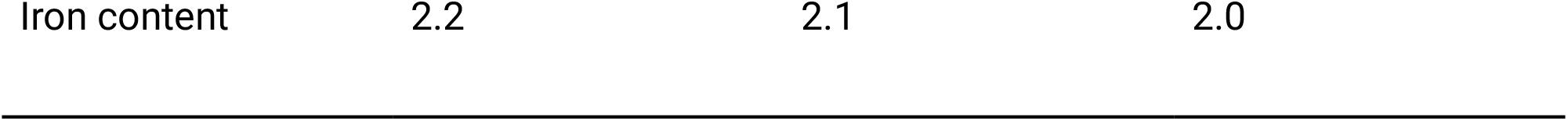
Percentage of Nutritional composition of *Pleurotus species* cultivated on Gmelina sawdust, Cassava peels and Oil palm fruit fibres respectively.

**Table 3:** Antioxidant values of Pleurotus species cultivated on Gmelina sawdust, Cassava peels and Oil palm fruit fibres respectively using DPPH assay procedure.

| Growth substrates used | Parameter |  |
| --- | --- | --- |
|  | RSA % | IC<sub>50</sub> mg/ml |
| Gmelina saw dust | 46 | 4.7 |
| Cassava peels | 74.9 | 2.4 |
| Oil palm fruit fibres | 61.6 | 3.1 |

Keys: RSA= Radical Scavenging Activity IC<sub>50</sub> = Maximum Inhibitory Concentration

**Figure 1:**
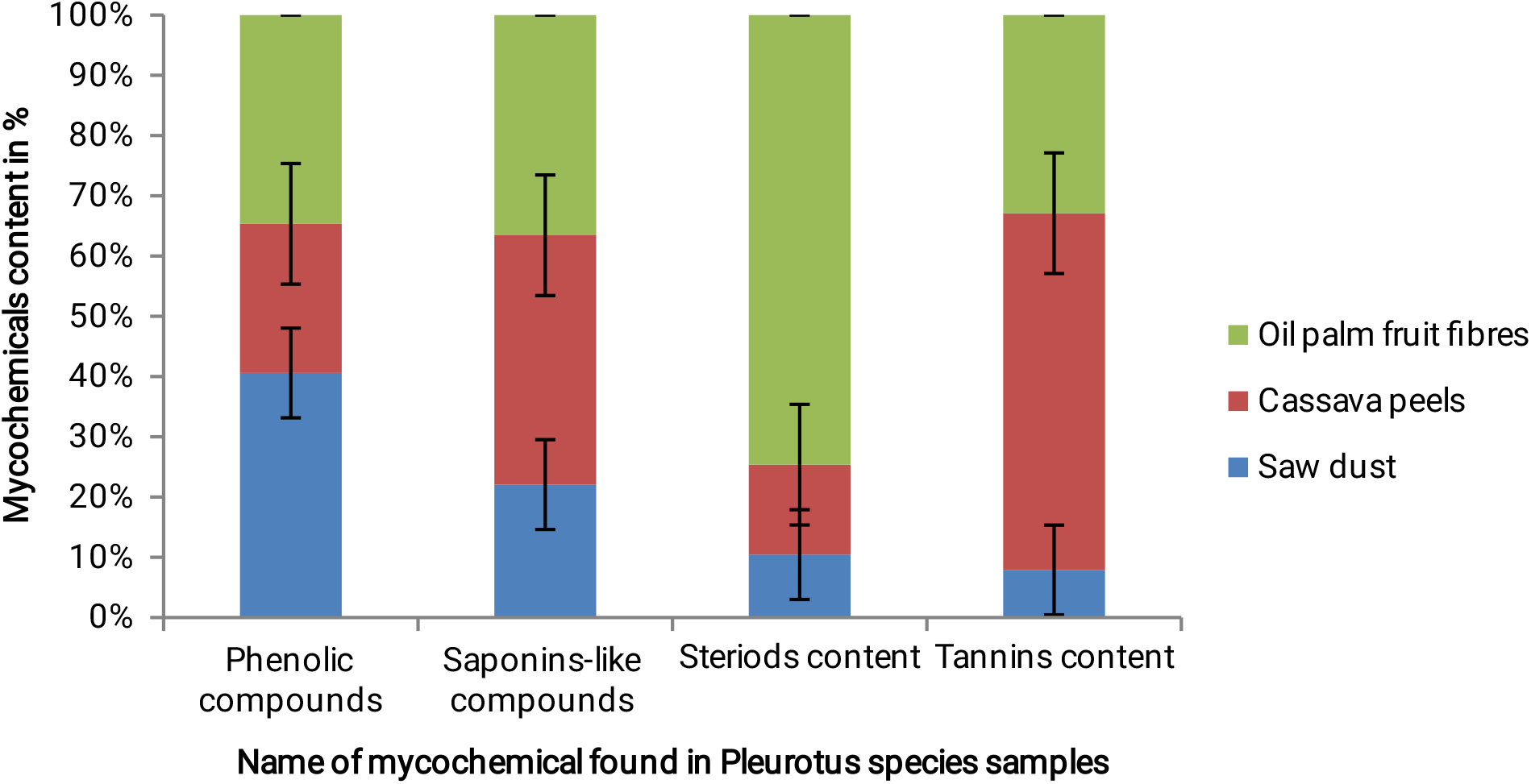
Percentage of mycochemicals in P. ostreatus cultivated on Gmelina sawdust, Cassava peels and Oil palm fruit fibres respectively

## 4.2. Discussion

The results of this study provide strong evidence that the selection of substrate plays a significant role in shaping the physical features, nutritional and bioactive properties of *Pleurotus ostreatus*, commonly known as oyster mushrooms. Specifically, the investigation reveals that the differences in the color and shape of *P. ostreatus* grown on different substrates used in the study may be due to differences in the minerals and organic compositions of the substrates, as also noted by Ogundele *et al*., 2017, and Ojha *et al*., 2025 in their findings. Also, elevated protein content observed in the mushrooms cultivated on Gmelina tree sawdust can be closely linked to the presence of nitrogen-rich compounds inherent in this particular substrate.These nitrogen compounds are instrumental in promoting enhanced protein synthesis during the growth and development of the mushrooms, thus contributing to their overall nutritional value. Hence, the composition and nature of the substrate used in growing mushrooms play a major role in determining the nutritional compositions and the mycochemicals present in such mushrooms. This finding closely aligns with the works of previous researchers (Fernandes *et al*., 2014; Akinrinola-Akinyemi *et al*., 2017; Mubashar *et al*., 2024;). Furthermore, the analysis indicates that the higher fiber content found in mushrooms grown on oil palm fruit fibers can be attributed to the naturally high fiber composition of the substrate itself. This characteristic not only reflects the substrate’s nutritional profile but also enhances the dietary benefits of the mushrooms, thereby potentially increasing their appeal as a healthful food source (Fakoya, 2025). Additionally, the study highlights that the increased antioxidant activity measured in mushrooms cultivated on cassava peels may be ascribed to the presence of various phenolic compounds within this substrate (Zahid *et al*., 2010: Alves et al., 2013; Pereira *et al*., 2013; Chang *et al*., 2018;). These phenolic compounds are known for their antioxidant properties, which can neutralize free radicals and contribute to overall health benefits (Wasser, 2015). As the concentrations of antioxidants increased, a noteworthy trend was observed in the study’s findings. Specifically, the IC_50_ values of the sample extracts exhibited an inverse relationship, indicating that as the concentration of antioxidants rose, the IC_50_ values correspondingly decreased. In practical terms, this means that higher concentrations of antioxidants were more effective in reducing the DPPH concentration, which is a common method used to measure antioxidant activity. This finding aligned closely with previous studies, notably the work conducted by Bakir *et al*. in 2018. Their research supports the notion that the strength of a compound’s antioxidant capacity can be quantified through its IC_50_ value. Furthermore, Pumtes *et al*. (2016) also highlighted this inverse correlation, noting that the IC_50_ value reflects the amount of antioxidant needed to achieve a 50% reduction in DPPH concentration. Thus, a lower IC_50_ value suggests a more potent antioxidant effect, as documented by Samruan *et al*. in 2012. As such, the substrate’s composition is critical in fostering these bioactive compounds, which may enhance the functional properties of *Pleurotus ostreatus*(Fakoya *et al*., 2014).

## 4.3. Conclusion

This study demonstrated that the substrate used for cultivation can significantly affect the nutritional content and bioactive compounds of *Pleurotus ostreatus*. By selecting suitable substrates, mushroom cultivators can produce mushrooms with specific desired characteristics, such as higher protein or fibre content, or enhanced antioxidant activity.

## 4.4. Recommendations

1. Based on the findings of this study, it is recommended that mushroom cultivators consider using Gmelina tree sawdust for producing high-protein mushrooms, palm fruit bunches fibres for producing high-fibre mushrooms, and cassava peels for producing mushrooms with enhanced antioxidant activity.
2. Future studies can investigate the effects of different substrate combinations on the nutritional content and bioactive compounds of *Pleurotus ostreatus*. Additionally, the potential health benefits of consuming mushrooms cultivated on different substrates can be explored.

## Disclaimer (Artificial intelligence)

Authors hereby declare that NO generative AI technologies such as Large Language Models (ChatGPT, COPILOT, etc.) and text-to-image generators have been used during the writing or editing of this manuscript.

## Authors declaration

**There is no conflict of interest among the authors**.

